# Identifying cell-to-cell variability in internalisation using flow cytometry

**DOI:** 10.1101/2021.11.24.469957

**Authors:** Alexander P Browning, Niloufar Ansari, Christopher Drovandi, Angus P R Johnston, Matthew J Simpson, Adrianne L Jenner

## Abstract

Biological heterogeneity is a primary contributor to the variation observed in experiments that probe dynamical processes, such as internalisation. Given that internalisation is a critical process by which many therapeutics and viruses reach their intracellular site of action, quantifying cell-to-cell variability in internalisation is of high biological interest. Yet, it is common for studies of internalisation to neglect cell-to-cell variability. We develop a simple mathematical model of internalisation that captures the dynamical behaviour, cell-to-cell variation, and extrinsic noise introduced by flow cytometry. We calibrate our model through a novel distribution-matching approximate Bayesian computation algorithm to flow cytometry data of internalisation of anti-transferrin receptor antibody in a human B-cell lymphoblastoid cell line. Our model reproduces experimental observations, identifies cell-to-cell variability in the internalisation and recycling rates, and, importantly, provides information relating to inferential uncertainty. Given that our approach is agnostic to sample size and signal-to-noise ratio, our modelling framework is broadly applicable to identify biological variability in single-cell data from internalisation assays and similar experiments that probe cellular dynamical processes.

## 1 Introduction

Endocytosis is the primary means by which cells uptake, or internalise, drugs, viruses, and nanoparticles [1–5]. Single-cell *in vitro* analysis of internalisation and similar dynamical processes reveals significant cell-to-cell variability in otherwise isogenic cell populations [6–12]. Such heterogeneity is ubiquitous to biology and essential to life, allowing for robust decision making, development, and adaptation of cell populations to environmental uncertainty [13–17]. From a clinical perspective, heterogeneity in drug uptake and response is considered a leading contributor to treatment variability and resistance [18, 19]. The challenges of working with data that comprises instrument noise and background fluorescence that often obfuscate biological variability means that it is relatively common for quantitative analysis of internalisation to neglect heterogeneity [20, 21]. Exacerbating these issues is a corresponding lack of mathematical tools that account for cell-to-cell variability and measurement noise while also providing information about the uncertainty in inferences and predictions drawn from noisy data.

Modern analysis technologies, including flow cytometry, allow the high-throughput collection of data from experiments that probe internalisation at rates exceeding a thousand cells per second (fig. 1) [22]. In an internalisation assay, material labelled with fluorescent probes are incubated with cells and internalised through the usual pathways (fig. 1a–b) [23, 24]. The fluorescence of surface-bound probes can be switched off by introducing a quencher dye, or that of internalised probes altered due to the lower pH in early endosomes [20, 23], providing quantitative information relating to the amount of material internalised. Flow cytometry provides measurements related to the total and internalised amount of material at various time points (fig. 1c–d). Since cells cannot otherwise be tracked between observation times, data comprise single-cell snapshots and individual trajectories are not available. While previous studies have shown that measurement noise introduced by the flow cytometry electronics and background autofluorescence are not insignificant; variability in the data is primarily biological [11, 25–29]. We confirm this by performing an internalisation assay with a dual-labelled fluorescent probe, finding that measurements are highly correlated, indicating a shared source of variability.

**Figure 1.**
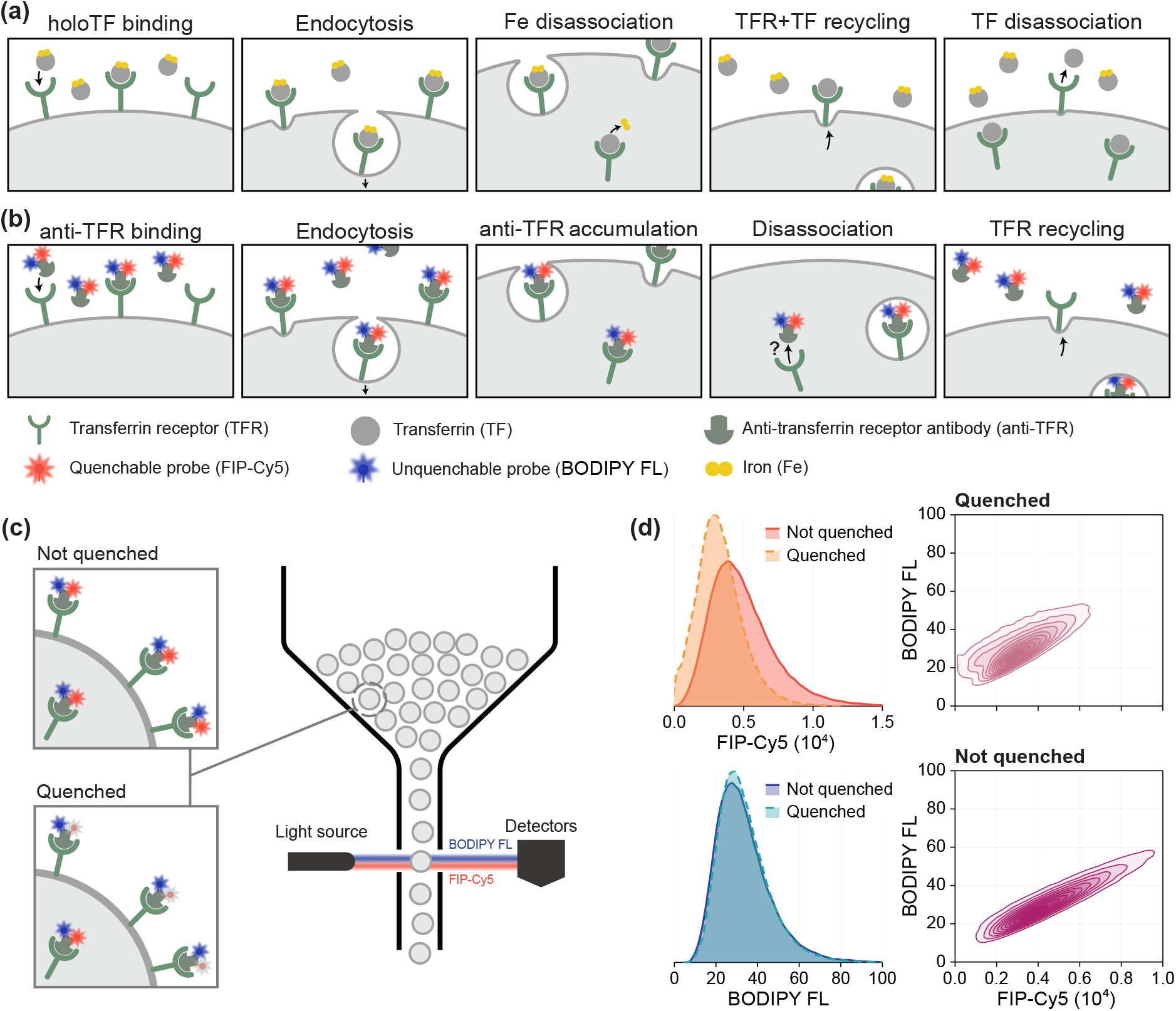
Internalisation dynamics and corresponding experimental assay. (a) Internalisation of transferrin, a protein responsible for the uptake of iron by cells. Iron-saturated transferrin (holoTF) binds to receptors on the cell surface and is internalised through *endocytosis*. In the low pH of endosomes, iron disassociates before the transferrin-receptor complex recycles to the cell surface. (b) A corresponding internalisation assay. Anti-transferrin receptor antibody (anti-TFR) dual-labelled with BODIPY FL and fluorescent internalisation probe (FIP)-Cy5 replaces iron-loaded transferrin and is internalised through the usual pathway. Experimental observations suggest that a small proportion of labelled antibody disassociates inside the cell, allowing receptor recycling and the accumulation of antibody inside the cell. (c) A quencher dye switches off fluorescence of surface-bound FIP-Cy5, providing information relating to the proportion of antibody that has internalised. Single-cell measurements of fluorescence from both probes is measured using flow cytometry. (d) Flow cytometry data obtained *t* = 10min after antibody are introduced. Since variability in the data is predominantly biological, data from each fluorescent label are highly correlated.

Mathematical and statistical techniques allow quantitative analysis of transient dynamics, heterogeneity, and measurement noise. As the number of molecules internalised by each cell is relatively large, single-cell trajectories describing the relative amount of material internalised can be accurately described by homogeneous models derived through kinetic rate equations. Ordinary differential equation (ODE) constrained Bayesian hierarchical and random effects models incorporate cell-to-cell variability through a parameter hierarchy where distributions parameterised by hyperparameters describe cell-level properties [30–32]. Both individual cell properties and hyperparameters are estimated during calibration of hierarchical models to data, presenting a significant computational challenge for the large sample sizes provided by flow cytometry data. In the mathematical literature, so-called heterogeneous [33] or random [34] ODEs and populations of models [35] make similar assumptions, often without assuming a parametric distribution of cell properties [9, 36, 37]. Issues presented by large sample sizes can be avoided by calibrating models using the empirical distribution of the data (through, for example, kernel density estimates) [33], an approach that provides point estimates but neglects inferential uncertainty and poses a challenge when the signal-to-noise ratio in the data is not sufficiently high.

In this study, we develop a mathematical model of internalisation that captures cell-to-cell variability by describing cell properties—specifically, the number of receptors, the internalisation rate, and the recycling rate of each cell—as jointly distributed random variables. To describe non-biological sources of variability from flow cytometry measurements of an internalisation assay, we couple the dynamical model to a probabilistic observation process that captures autofluorescence and measurement noise. We take a Bayesian approach to parameter estimation and develop a novel approximate Bayesian computation (ABC) [38–40] algorithm that matches distributional information from flow cytometry measurements. This approach is agnostic to sample size and the signal-to-noise ratio and provides point parameter estimates and information relating to inferential uncertainty.

We demonstrate our approach by studying heterogeneity in the internalisation of anti-transferrin receptor (anti-TFR) antibody in C1R cells, a human B lymphoblastoid line. Data comprise potentially noisy flow cytometry measurements from an internalisation assay developed in our previous work, specific hybridisation internalisation probe (SHIP) (fig. 1b–c) [20,41]. Measurements are collected from anti-TFR antibody dual labelled with BODIPY FL and fluorescent internalisation probe (FIP)-Cy5. We take measurements both with and without a quencher dye, which only switches off the fluorescence of surface-bound FIP-Cy5 without affecting the BODIPY FL signal or internalised FIP-Cy5. Therefore, we obtain jointly distributed data that comprises noisy measurements of the total and internalised amount of antibody in each cell (fig. 1c–d). Snapshots are collected from samples that are incubated with antibody saturated medium for various periods of time to provide measurements relating to both the total and internalised amounts of antibody present on each cell. Using our mathematical model, we are able to identify key sources of biological variability and provide predictions that give insight into how the uptake of material varies between cells. Importantly, our approach to parameter inference enables us to quantify the uncertainty in inferences made, allowing us to provide experimental design guidance.

## 2 Results

### Dynamical model of internalisation

We describe the internalisation of antibody and the recycling of receptors using a compartment model. Given the concentration of antibody in the surrounding medium is sufficiently high, we assume that the association rate of antibody to free receptors on the cell surface is much higher than the kinetic rates of internalisation and recycling (supplementary material, S1). Therefore, we describe the number of antibody-receptor complexes on the cell surface, *S*, and that endocytosed, *E*. Before incubation in antibody-saturated medium, *T* endocytosed receptors are bound to transferrin. In the experiments, we observe that the total association of antibody with cells continues to increase, consistent with receptor recycling and accumulation of fluorescent material inside the cell, which we capture by describing the number of free endocytosed antibody, *F*. These assumptions give rise to the dynamical model (fig. 2a)

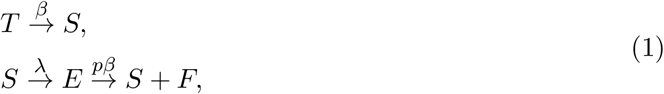

where *β* [min^−1^] is the recycling rate, *λ* [min^−1^] is the internalisation rate, and *p* is the probability that an endocytosed antibody disassociates allowing receptor recycling. It is also possible that endocytosed antibody, *E*, can return to the cell surface without disassociation from the receptor. However, we have not included this in our model as a recycled antibody-receptor complex is indistinguishable from that bound on the cell surface, *S*.

**Figure 2.**
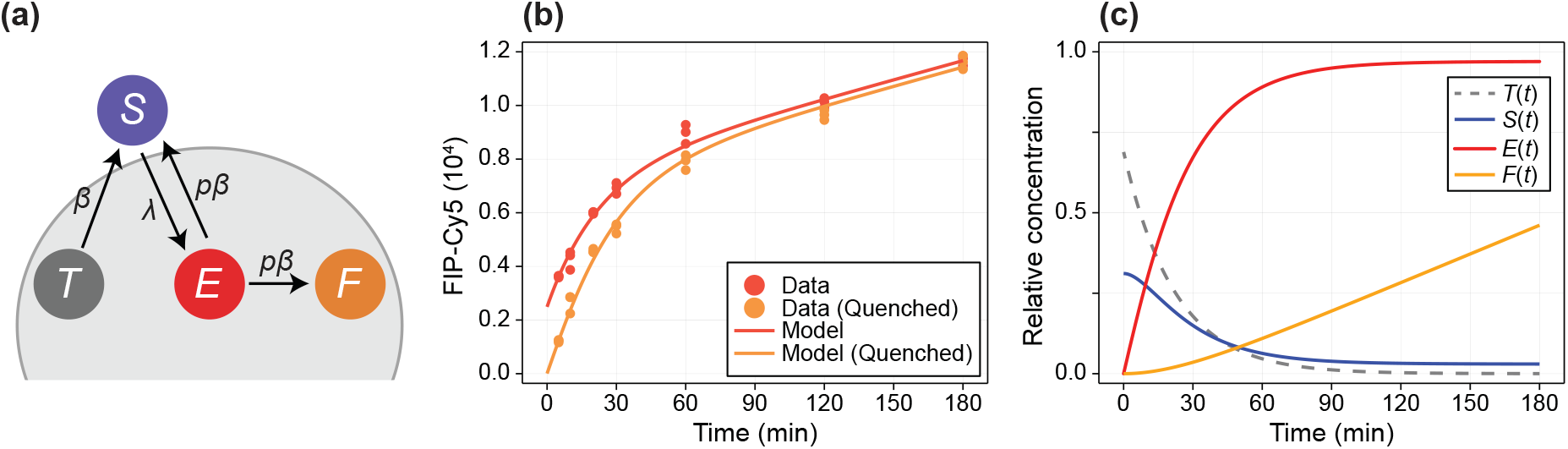
Dynamical model of internalisation and recycling matches experimental data. (a) The dynamical model describes the relative concentration of internal, transferrin bound receptors, *T* (grey); surface antibody-bound receptors, *S* (blue); internal antibody bound receptors, *E* (red); and internal free antibody, *F* (orange). (b) MFI of FIP-Cy5 fluorescence measurements for samples that are not quenched (red) and those that are (orange) at various time points. The dynamical model is calibrated using maximum likelihood estimation, with the solution shown (solid curve). (c) Solution to the mathematical model (Methods, eq. (M16)) at the maximum likelihood estimate (table 1).

Given that the number of receptors in each cell is relatively large, eq. (1) can be formulated as a linear ordinary differential equation with exact solution **x**(*t*) = **x**_0_ exp(**M***t*), where **M** is a matrix of coefficients and **x**(*t*) denotes the number of molecules in each compartment (Methods M.2.2). Initially the system is in equilibrium, so

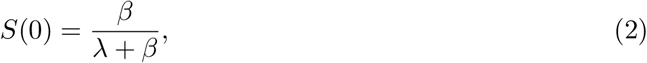

where molecule counts are taken with respect to the total number of receptors on the cell, denoted *R*, so *S*(0) + *T*(0) = 1.

### Inference using mean fluorescence intensity measurements

Flow cytometry measurements are typically summarised using mean fluorescent intensity (MFI); point statistics equivalent to the geometric mean of the fluorescence distribution (fig. 2b). Cy5 MFI measurements from samples that are not quenched are related to the total amount of antibody in the sample, *A*(*t*) = *S*(*t*) + *E*(*t*) + *F*(*t*), and measurements from quenched samples are related to the amount of internal antibody in the sample, *I*(*t*) = *E*(*t*) + *F*(*t*). Assuming incomplete quenching so the probability that a Cy5-labelled probe is quenched is *η* ≈ 0.94 (supplementary material, S2) and average autofluorescence *E_Q_*, MFI measurements can be modelled by

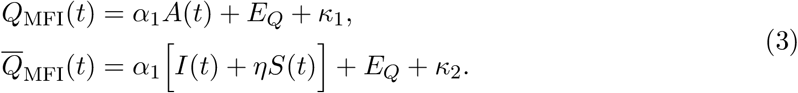

Here, we denote by *Q*_MFI_(*t*) MFI measurements from the FIP-Cy5 (i.e., quenchable) probe in the samples that are not quenched, and by 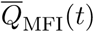 that of quenched samples. We capture variability in MFI measurements between experiments by assuming measurement error *κ*_1_, *κ*_2_ ~ Normal(0, *σ*^2^). We refer to eq. (3) as the *homogeneous model* since the dynamical parameters *λ* and *β*, and the number of receptors, *R*, does not vary cell-to-cell and are fixed for the population.

To assess the suitability of the dynamical model and provide a baseline to assess our model that captures biological heterogeneity, we calibrate eq. (3) to experimental data using maximum likelihood estimation. We tabulate estimates and confidence intervals approximated using the observed Fisher information in table 1, and show the model best-fit in fig. 2b.

**Table 1.**
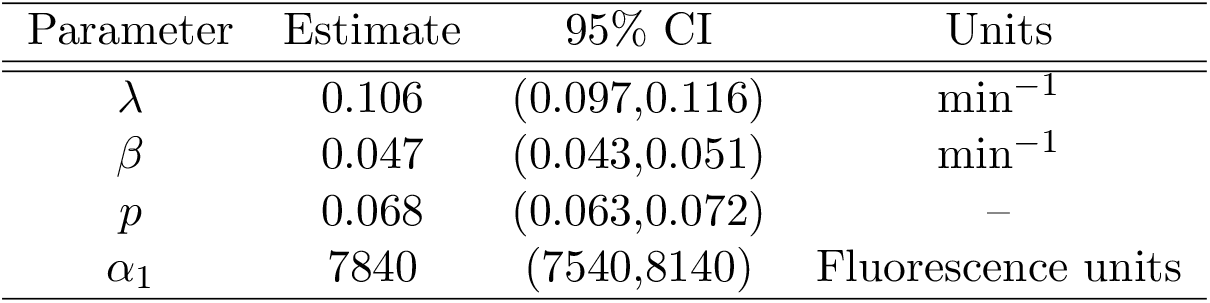
Parameter estimates and approximate confidence intervals for the homogeneous model. Approximate confidence intervals are calculated using the observed Fisher information matrix, calculated from the Hessian of the log-likelihood function [42].

The homogeneous model provides a fit that qualitatively matches MFI measurements from the experimental data (fig. 2b), and all parameters are identifiable within relatively precise intervals (table 1). Estimates for the internalisation and recycling rates suggest that a proportion of approximately

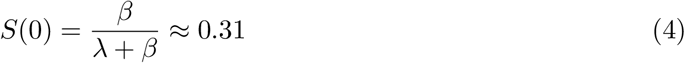

of transferrin receptors lie on the surface at equilibrium. Estimates for *p* suggest that 6.8% (95% CI (6.3%,7.2%)) of internalised antibody disassociates, allowing receptor recycling. This is also evident from simple observations of the experimental data, since the fluorescent intensity increases throughout the experiment, suggesting that a small proportion of receptors remain on the surface while antibody accumulates inside the cell (fig. 2c).

### Incorporating biological variability into dynamical model of internalisation

We assume biological variability arises through both physical and physiological differences between cells in the population. Specifically, we allow number of receptors, *R*, and dynamical parameters *λ* and *η* to vary cell-to-cell. Without loss of generality, we set 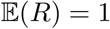 so receptor and antibody counts are taken with respect to the average receptor count in the population.

The properties of the *i*th cell are given by the random variable ***ξ**_i_* = (*R_i_, λ_i_, β_i_*). We assume unimodality and a parametric description of ***ξ**_i_* that allows us to estimate the first three moments and dependence structure. Marginals are given by

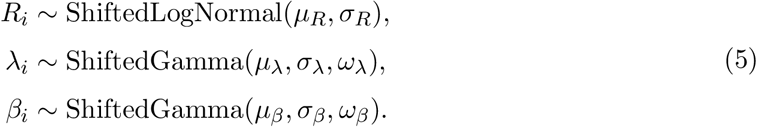

where *λ_i_* and *β_i_* are shifted Gamma variables parameterised in terms of their respective means, standard deviations and skewnesses (supplementary material, S4). The number of receptors is assumed to be shifted log-normally distributed [43]. To ensure positivity, we truncate ***ξ**_i_* so that *β_i_, λ_i_, β_i_* ≥ 0. We model the dependence structure of ***ξ**_i_* with a Gaussian copula parameterised by the correlation matrix

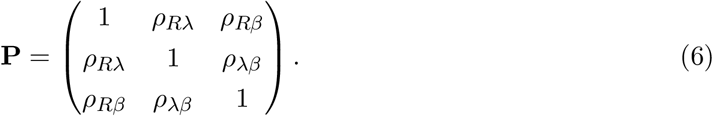

To ensure **P** remain positive definite, we infer *ρ_Rλ_*, *ρ_λβ_* and 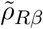 (all constrained to the interval (−1, 1)) where

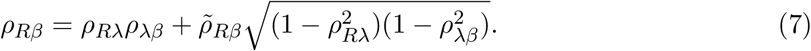

Therefore, *ρ_Rλ_* (and similarly for *ρ_Rβ_* and *ρ_βλ_*) describes the strength of the correlation between the number of receptors, *R*, and internalisation rate, *λ*.

The *heterogeneous model* is a random ODE model where **x**(*t*) and its constituents are random variables [34]. For example, *A*(*t*) is a random variable describing the distribution of bound-antibody a cell at time *t*.

### Statistical model for flow cytometry data

Measurement noise in flow cytometry is primarily attributable to shot noise introduced from the photomultiplier tubes (PMT noise) that convert the photon signal to an amplified, analogue electrical signal. Recent studies suggest that the square coefficient of variation of such noise is approximately constant [28], so we model shot noise with uncorrelated white noise (i.e. Gaussian), with variance proportional to the true signal. The second source of noise is cellular autofluorescence, where the laser used to excite the labelled antibody can excite other molecules in the cell, leading to a background autofluorescence where signal is present in the absence of antibody. We build an empirical distribution of autofluorescence (*E_Q_, E_U_*) using a sample where cells have not been introduced to labelled antibody (supplementary material, S3).

We interpret measurements from the FIP-Cy5 probe, which is quenchable and denoted by *Q*(*t*), and the BODIPY FL fluorescent dye, which is not quenchable and is denoted by *U*(*t*). Therefore,

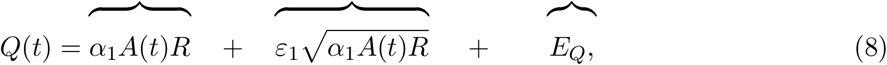

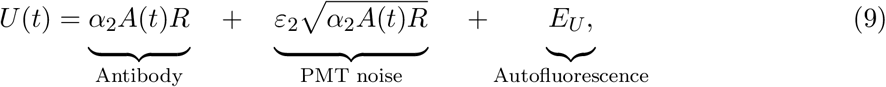

where 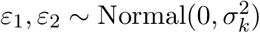. Similarly, the measurements from quenched samples are given by

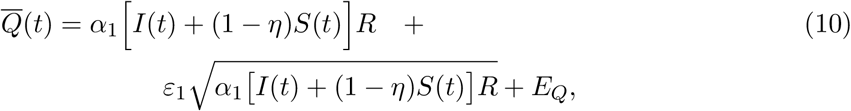

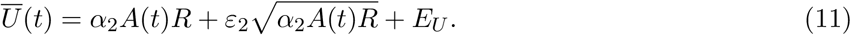

### Calibration and uncertainty quantification

We take a Bayesian approach to parameter estimation, calibrating the noisy heterogeneous model to SHIP assay data using a novel approximate Bayesian computation (ABC) algorithm that matches the empirical distribution of flow cytometry measurements, under the assumption that measurements from each probe are linearly correlated (fig. 3a–b). Model simulations based on *n* = 1000 cells per observation time, per condition (quenched or not quenched) are compared to experimental data using the Anderson-Darling distance [44] and Pearson correlation. Before observation of data, knowledge about the parameters are encoded in prior distributions, chosen to be independent and uniform with bounds given as axis limits in fig. 3. Full details are given in Methods M.2.3.

**Figure 3.**
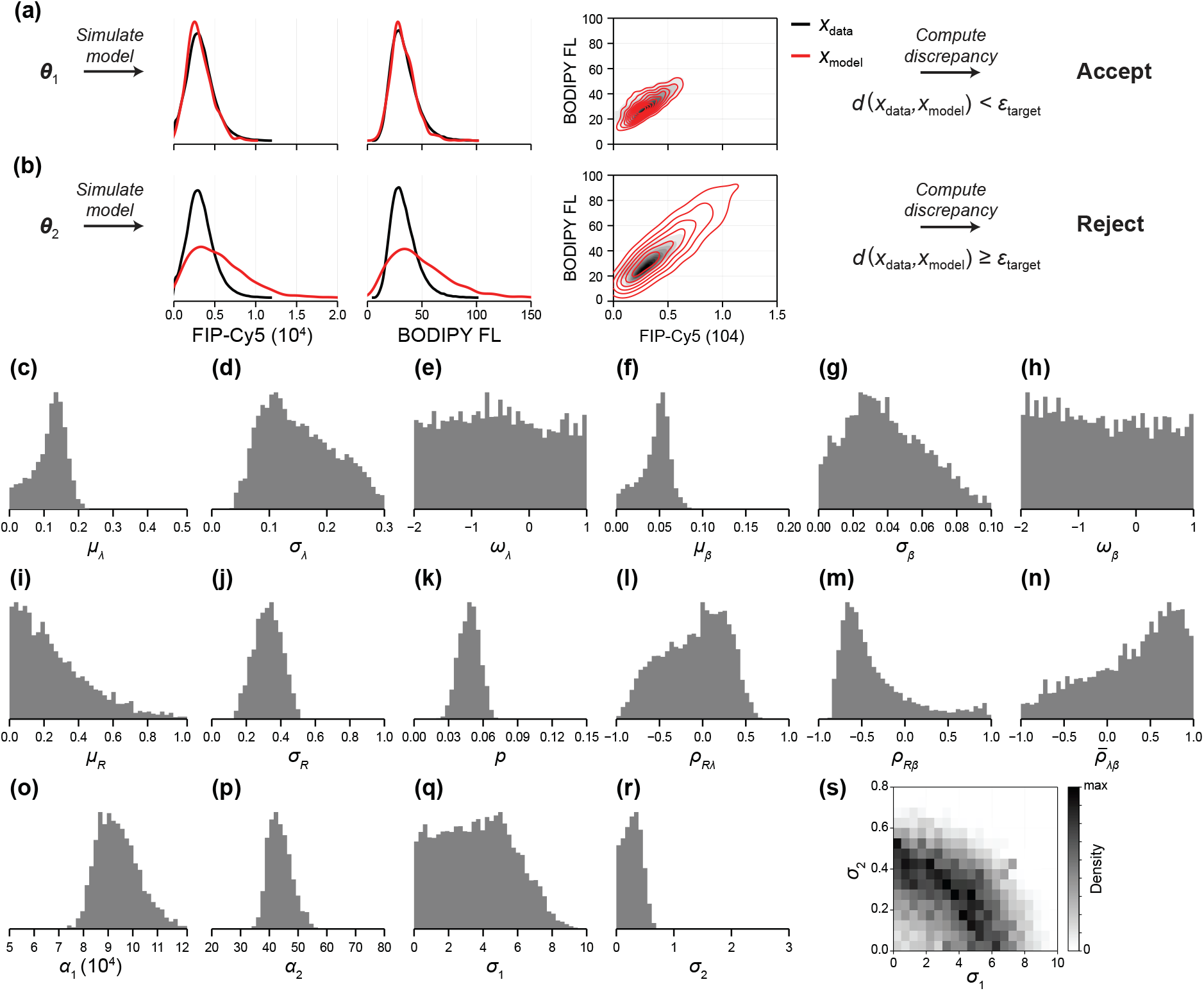
Model calibration and uncertainty quantification using ABC MCMC. (a–b) In ABC, data are compared to model simulations using a weighted sum of Anderson Darling distances and discrepancy in the correlations. (a) Parameter combinations that produce model realisations sufficiently similar to the experimental data, i.e. ***θ***_1_, are accepted as posterior samples. (b) Parameter combinations that do not, i.e. ***θ***_2_ are rejected. (c–s) *Posterior samples* obtained using ABCMCMC represent parameter combinations that produce realisations of the model that are similar to experimental observations. Chains are initiated at the global minimum identified using ABC SMC, and every 100th sample is retained. All axis limits for univariate distributions correspond to the prior support (uniform priors are used). Parameter descriptions and MCMC diagnostics are given as supplementary material (S5).

First, we employ a particle filter based on sequential Monte Carlo (SMC) [45] to identify the region of the 16-dimensional parameter space that produces model realisations that lie close to the data and to establish a threshold below which model simulations are deemed sufficiently close to the experimental data. We then apply a Markov chain Monte Carlo (MCMC) algorithm to explore the support of the posterior distribution, interpreted as the region of the parameter space that produces model simulations that match experimental data, quantifying uncertainty in parameter estimates. In fig. 3 we plot posterior samples from four independent tuned MCMC chains thinned to a total of 400,000 samples, providing an effective sample size of at least 1,000 per parameter. To visualise model predictions, we compute a point estimate by further thinning the chains to a total of 400 samples, and identifying the parameter set that produces the lowest average discrepancy from 100 model realisations. Model predictions at the point estimate are shown alongside experimental data in fig. 4. MCMC diagnostics, parameter descriptions and best-fit estimates are given in supplementary material (S5).

**Figure 4.**
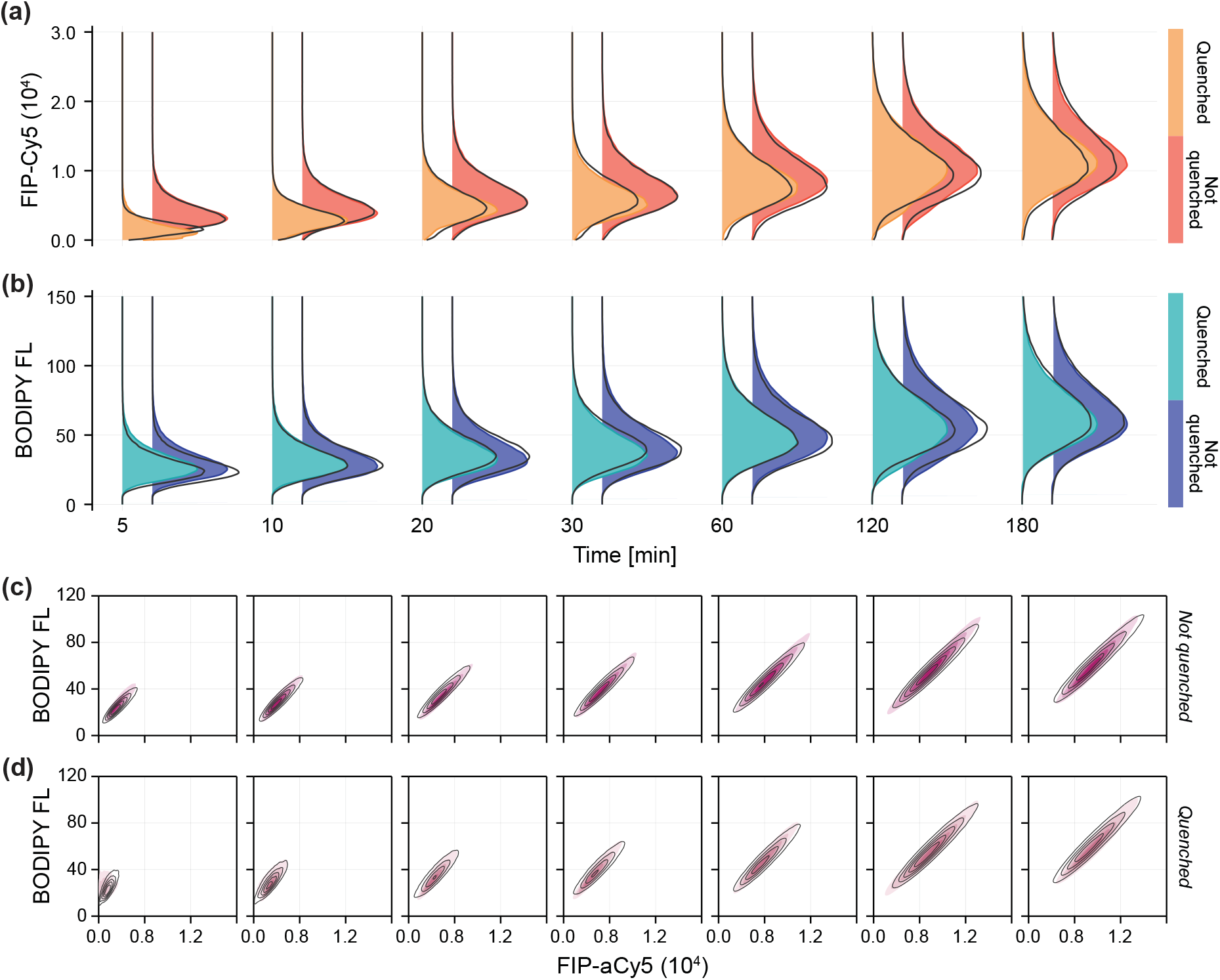
Mathematical model captures variability in experimental data. (a–b) Univariate kernel density estimate of the fluorescent intensity distribution from (a) the Cy5 probe, which is susceptible to the quencher dye, and (b) BODIPY FL, which is not. In each case, the distribution from the quenched experiment is shown to the left in the lighter colour. (c–d) Bivariate kernel density estimates of the joint fluorescent intensity distributions for FIP-Cy5 and BODIPY FL measurements. The model prediction from a synthetic data set of 100, 000 cells per observation time, per condition, is overlaid in black.

### Heterogenous model captures biological variability

The heterogeneous model produces realisations with excellent agreement to flow cytometry measurements, matching both marginal and jointly distributed data from both probes (fig. 4). Minor discrepancies in univariate distributions highlight the main sources of unaccounted error; for example, error relating to the precise time at which internalisation is ceased and error relating to flow cytometry gating.

Samples relating to the skewness of the internalisation and recycling rate distributions, *ω_λ_* and *ω_β_*, respectively, (fig. 3c,f) show that information in the experimental data is insufficient to identify the shape of the internalisation and recycling rate distributions. While the precision to which we can identify the variance of each rate, *σ_λ_* and *σ_β_*, (fig. 3b,e) is limited, it is clear that *σ_λ_* > 0.057 (lower bound on a 95% CrI), providing evidence to suggest heterogeneity in the internalisation rate. While *p* has the same interpretation between the heterogeneous and homogeneous models, the estimates from the heterogeneous model, *p* = 4.7% (95% CrI (3.0%,6.5%)), are lower with a greater amount of uncertainty than in the homogeneous model.

In fig. 5 we plot the inferred distributions of *R*, *λ* and *β*. To visualise uncertainty in estimates of these distributions, we show a 95% credible internal (CrI) for the univariate probability density functions by resampling from the posterior distribution. Compared to distributions of the dynamical parameters *λ* and *β*, the distribution of the relative receptor count, *R*, is identified with much greater precision (fig. 5a). *R* does not feature in the dynamical model and is, therefore, less sensitive to issues relating to model misspecification. While results in fig. 3j–l show relatively large uncertainty in the correlations between parameters, it appears likely that the receptor count and recycling rate are negatively correlated (83% of posterior samples have *ρ_Rβ_* < 0). Parameters identified in the homogeneous model based on MFI measurements are contained within high density regions of the inferred distributions in the heterogeneous model. This is also the case when estimates are compared to bivariate distributions in fig. 5d–f; however, the interpretation of the homogeneous model parameters in the context of data with significant heterogeneity is unclear, highlighting the importance of modelling biological variability when interpreting flow cytometry data of dynamical processes like internalisation.

**Figure 5.**
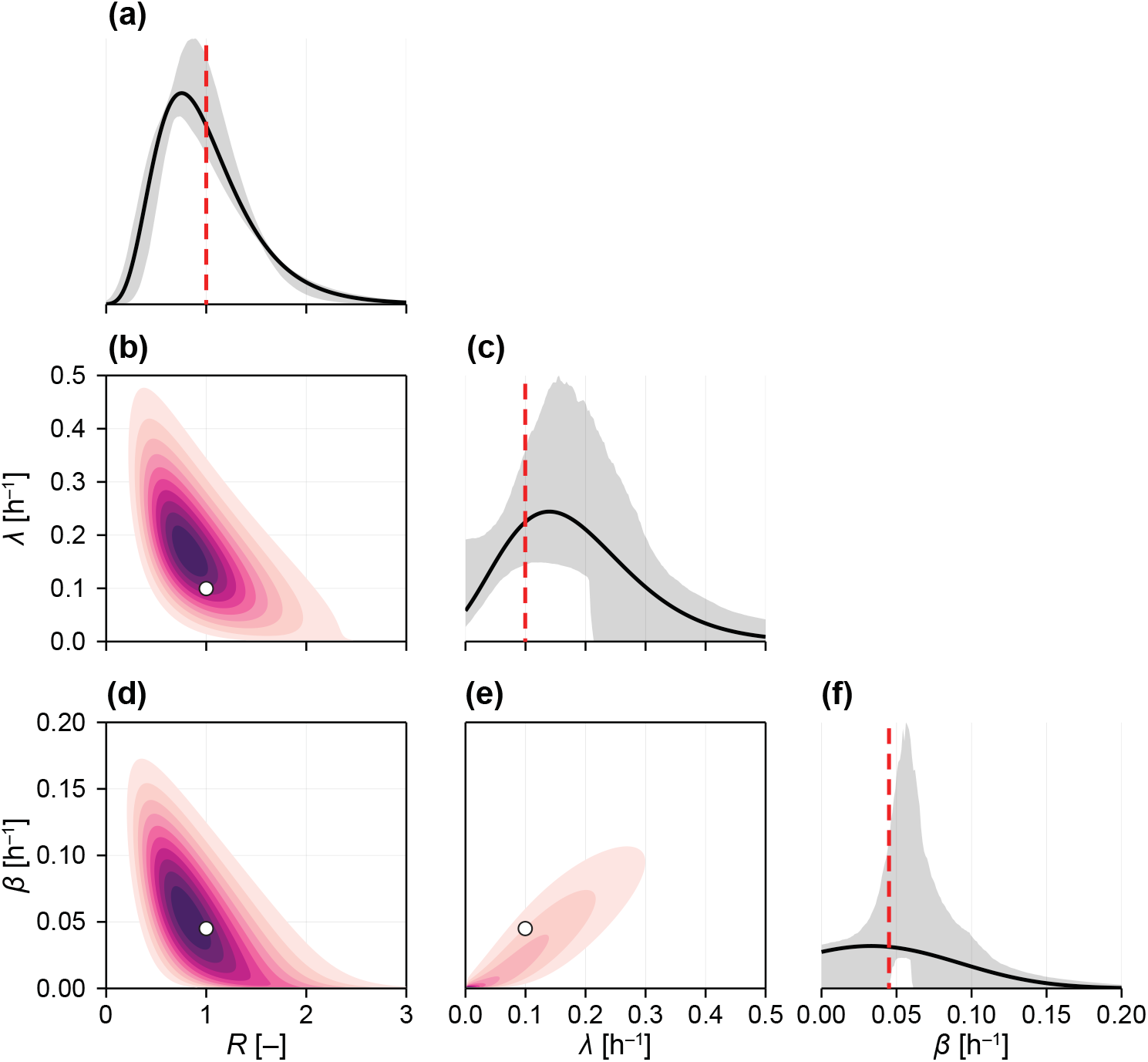
Inferred parameter distributions and associated uncertainty. Inferred distribution of (a) the relative number of receptors, *R*; (c) the internalisation rate, *λ*; and, (f) the recycling rate, *β*. Shown are the distributions at the best-fit (black), a 95% credible interval of the respective probability density functions constructed from re-sampled MCMC samples (grey), and estimates from the homogeneous model (red). (b,d,e) Bivariate distributions at the best-fit. Estimates from the homogeneous model are shown in white.

The data appears insufficient to distinguish between PMT noise from each fluorescent channel. Initial examination of estimates for the relative magnitude of the noise from the quenchable (FIP-Cy5) and unquenchable (BODIPY FL) probe signals in fig. 3o–p suggests that the nonoise model may be appropriate, lending the study to analysis of models that assume negligible noise [33]. However, the joint distribution of *σ*_1_ and *σ*_2_ (fig. 3s) reveals an elliptical region, suggesting that the model requires PMT noise in the signal from *at least one* probe. Similar phenomena are observed in error-in-variables or total-least-squares problems, where errors are introduced in both independent and dependent variables, and only the ratio of the error variances is identifiable [46].

### Model predicts unobservable measurements

A primary goal of flow cytometry analysis is to quantify the amount of fluorescent material present in a sample. In the context of an internalisation assay, we are interested in the *proportion* of material internalised through time. By accounting for variability introduced through receptor count, PMT noise and autofluorescence, we are able to better quantify the amount, or proportion internalised, of antibody compared with standard approaches.

Since it is not possible to collect noise-free data relating to the joint distribution of *I*(*t*), provided from quenched samples, and *A*(*t*), provided from sampled that are not quenched, statistics such as the proportion of antibody internalised by each cell cannot be directly measured. Rather, such statistics are typically estimated as

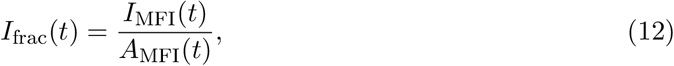

where *I*_MFI_(*t*) and *A*_MFI_(*t*) are scalar estimates of the average proportion of internal and total antibody estimated using MFI [7]. Using our calibrated heterogeneous model, we can predict the *distribution* of material internalised through time by simulating the model with sources of noise removed. In fig. 6a, we show the time-evolution of the distribution of *I*(*t*)/*A*(*t*) at the model-best-fit, along with the equivalent prediction from the homogeneous model. In fig. 6b, we repeat this exercise for the total relative amount of antibody internalised, *I*(*t*). To understand uncertainty in these distributions, in fig. 6c we show the time evolution of the distribution of *I*(*t*)/*A*(*t*) alongside credible intervals formed by resampling parameters from the posterior distribution.

**Figure 6.**
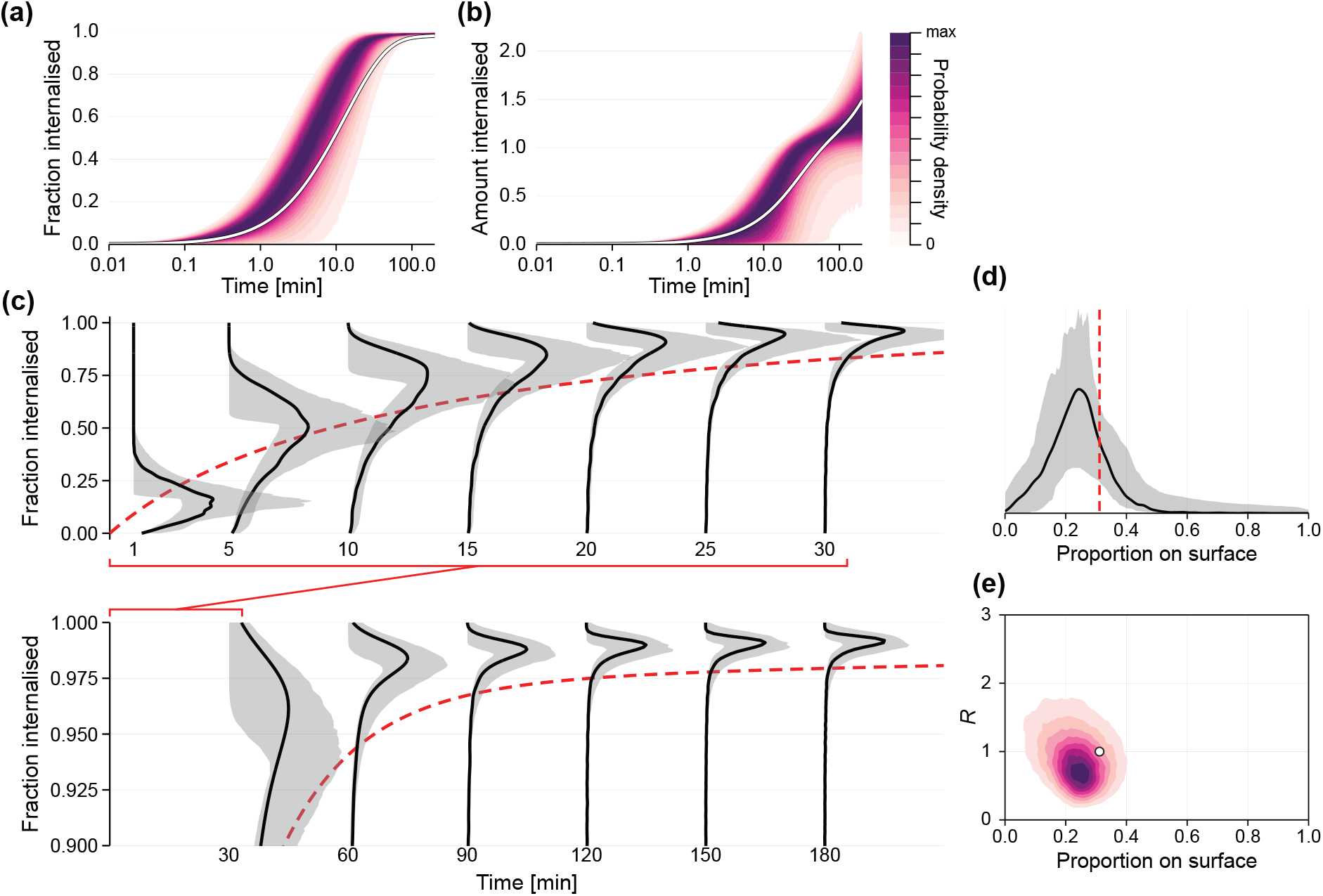
Model predicts unobservable measurements. Using the calibrated mathematical model, we can predict the time-evolution of the distribution of (a,c) fraction antibody inside the cell; (b) amount of antibody internalised relative to the average number of receptors on a cell, *I*(*t*); and (d–e) proportion of receptors on the cell surface at equilibrium. (a,b,e) Show the distribution at the model best-fit, which is shown in black in all other plots; (c,d) additionally show 95% credible intervals constructed from MCMC samples. (c) To compare predictions with the homogeneous model, we show *A*(*t*)/*I*(*t*) predicted by the homogeneous model (red-dashed) using only MFI measurements. In (a,b) distributions are normalised to the mode.

While our analysis revealed that several parameters are non-identifiable, or cannot be constrained to a relative precise interval, we are still able to produce relatively precise predictions of statistics such as the proportion of material internalised. Results in fig. 6c show a discrep-ancy between predictions from the homogeneous and heterogeneous models. Aside from very early time, when the distribution of material internalised is relatively wide, the homogeneous model predictions lie within the lower tail of the predicted distribution. This is consistent with the discrepancy we observe in estimates of *p* between models: the heterogeneous model predicts that antibody disassociation and receptor recycling is rarer that what is predicted by the homogeneous model. This results in a smaller proportion of surface-bound antibody at late time.

Using the inferred joint distribution of *λ* and *β* we can build a picture of the proportion of receptors present on the cell surface at equilibrium (i.e., at the start of the experiment), *S*(0) (eq. (4)). In fig. 6d, we show the inferred distribution of *S*(0) at the model-best-fit, along with the uncertainty associated with the estimate and that predicted by the homogeneous model. While we have not precisely estimated this distribution, it is clear that, on average, a smaller number of receptors are present on the cell surface than not, in agreement with the prediction of the homogeneous model of 31%. We also see that the inferred distribution is highly variable; at the best-fit, for example, non-zero density at zero suggests that some cells have a very small proportion of receptors on the surface, perhaps due to inhibition of recycling. In fig. 6e, we show the inferred relationship between *S*(0) and *R* at the best-fit. This result suggests that cells with a larger number of receptors—which may correlate to cells in latter stages of the cell cycle—have fewer surface-bound receptors.

## 3 Discussion

Heterogeneity is ubiquitous in cell processes such as the internalisation of material, yet the phenomenon is poorly understood and often ignored. Paired with experimental protocols that probe these processes, flow cytometry is capable of generating vast quantities of single-cell-snapshot data that captures cell-to-cell variability. Often, such data are summarised with point statistics that provide information about the transient behaviour to the detriment of acknowledging variability between otherwise isogenic cells. In this study, we develop a mathematical model of internalisation that captures dynamical behaviour, biological variability, and measurement noise of arbitrary magnitude. We apply our model to identify key sources of biological variability in the internalisation of anti-TFR antibody by C1R B-cell lymphoblastoid cells.

While computationally costly, our distribution-matching ABC approach to inference carries several advantages over likelihood-based approaches; for example, those based on Bayesian hierarchical models or those that model cell properties as a finite mixture [37]. First, ABC is robust to model error, incorporating uncertainty due to factors that are not explicitly modelled [47]. This might include the relatively small discrepancies we observe in fig. 4 that highlight potential model-misspecification as well as error introduced experimentally, such as the precise measurement time and the time at which internalisation is ceased.

Secondly, the distribution-matching approach allows the interpretation of pre-processed or summarised data. Automatic clustering algorithms [48–50] are fast replacing manual gating, providing an opportunity to analyse the parametric mixture distributions identified algorithmically, rather than relying on accurate classification of individual data points to perform analysis on the underlying data. Lastly, our approach is agnostic to both the size of the dataset and the complexity of the underlying measurement model. While the signal-to-noise ratio in our data is high (demonstrated by the amount of variability we identify as biological in origin), this is not always the case. In particular, flow cytometry measurements are often corrupted by autofluorescence and bleed-through from overlapping emission spectra. In our framework, both sources of extrinsic variability can be built into the probabilistic observation process or accounted for using pre-processing software where the compensated distributions are analysed rather than the underlying data.

Working with single-cell snapshot data provides limited information compared to single-cell trajectory data, potentially explaining why inferences relating to heterogeneity in dynamical parameters are relatively imprecise (fig. 5). In particular, the data provides no information about the joint distribution of antibody concentration between observation times. Additional results in fig. 7 illustrate the predicted dependence in internalised antibody concentration between early (10min) and late (120min) observation times, denoted by *Q*(10) and *Q*(120), respectively. An interpretation of model predictions with higher fitted correlations between *Q*(10) and *Q*(120) is that single-cell trajectories remain ordered: cells with a relatively lower proportion of antibody internalised at *t* = 10min retain a relatively lower proportion at *t* = 120min. Therefore, it is unclear from the available data (fig. 7b) whether cell trajectories remain ordered or whether cells can “catch up”; that is, whether cells that are initially slow to internalise material end up with a large amount internalised at later time points. Intuitively, assuming that such a correlation is strong (i.e., trajectories remain ordered) strongly impacts inferences. Our results in fig. 7c–d show that making such an assumption narrows uncertainty in the distribution of recycling rates to distributions where cells that do not recycle (i.e., *β_i_* = 0) are rare.

**Figure 7.**
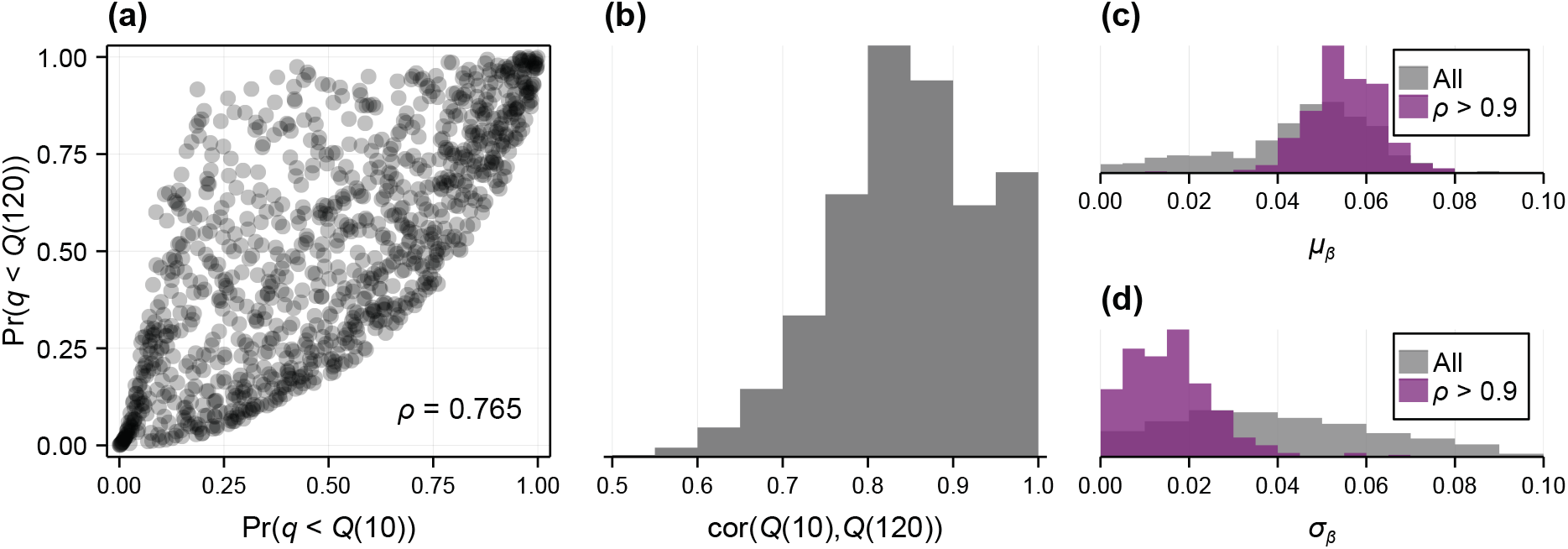
Dependence between the amount of material internalised at successive observation times remains uncertain. We demonstrate model predictions and uncertainty in the dependence between the noise-free quenched signal at time *t* = 10 and *t* = 120 min. (a) Quantile-quantile plot for a simulation with 1000 cells at the best-fit. Correlation of *ρ* = 0.765 is calculated based on a fitted Gaussian copula. (b) Uncertainty in the inferred correlation based on resampled posterior samples. (c,d) Assuming strong dependence between observation times affects inferences. We show the posterior distribution for the recycling rate mean and standard deviation parameters, *μ_β_* and *σ_β_*, respectively, if we assume a correlation of at least 0.9.

Aside from stochastic variations between isogenic cells—due to gene expression [51], for example—variability in internalisation is at least partially driven by the cell cycle [52, 53]. Therefore, we might expect lower internalisation and recycling rates in cells preparing to undergo mitosis which, therefore, have a larger number of receptors. This is also suggested by results relating to the best-fit in fig. 5, which show that the internalisation and recycling rates decrease with the number of receptors. These results raise the possibility of a non-Gaussian dependence between dynamical rates that our model cannot capture. For example, the dependence between *R* and *λ* may not be monotonic: internalisation by cells in very late stages of the cell cycle might be inhibited, whereas in general, larger cells may internalise material more quickly [54]. Distribution-free approaches [37] might better capture the dependence structures in these cases. However, given that our model is already able to match the experimental data, adding complexity will exacerbate parameter non-identifiability. Therefore, further work should focus on experimental design [55]; by inhibiting recycling, or pre-sorting cells to remove variability in *R*, for example.

Our analysis demonstrates that inferences drawn using approaches that neglect heterogeneity can be misleading. In particular, the interpretation of predictions and parameter estimates from the homogeneous model is mathematical unclear. Generally, realisations of the homogeneous model do not represent the mean of realisations of the heterogeneous model, nor do they represent realisations where parameters in the heterogeneous model are first averaged [34]. While, in our case, parameters identified by the homogeneous model are contained within the distribution identified by the heterogeneous model, the homogeneous model produces biased predictions that are not representative of the entire population (fig. 6). These findings highlight a need to co-develop mathematical tools that account for biological variability in analysis of single-cell data.

A better understanding of heterogeneity in internalisation has important implications for drug delivery [5,19], in addition to our understanding of pathological processes, such as the internalisation of viruses [56,57]. In this study, we develop a novel quantitative model that captures biological variability in internalisation using arbitrarily noisy flow cytometry data. In contrast to conventional approaches, we can produce predictions that give insight into the variability in material internalised while accounting for inferential uncertainty. Applying mathematical models that capture biological variability allows practitioners to get the most out of the vast amounts of single-cell data generated by flow cytometry and other modern experimental tools.

## Data availability

Code and data are available on GitHub at github.com/ap-browning/internalisation.

## Author Contributions

All authors interpreted the results and conceived the study. A.P.B. performed the mathematical analysis and implemented computational algorithms. N.A. performed the experiments. A.P.B. and N.A. drafted the manuscript. All authors provided comments and gave approval for publication.

## M Methods

### M.1 Experimental methods

#### M.1.1 Cell culture

C1R cells, a human B cell lymphoblast cell line, were cultured in Dulbeccos Modified Eagle Medium (DMEM) supplemented with 10% FBS and 1% Penicillin Streptomycin, at 37°C in a humidified 5% CO_2_ atmosphere.

#### M.1.2 Dual-labelled fluorescent internalisation probe

Purified monoclonal IgG1 anti-human transferrin receptor antibody (OKT9) [58] was purchased from WEHI Antibody Facility.

The antibody was labelled with two fluorescent dyes; BODIPY FL and FIP-Cy5. For this, anti-TFR antibody was incubated with BODIPY FL-NHS ester and incubated at 4 °C overnight. BDP FL-labelled antibody was purified using a 7KMWCO Zeba spin desalting column (Thermo Scientific). The antibody was then functionalised with dibenzylcyclooctyne (DBCO)-NHS ester. Functionalised antibody was purified using a 7K MWCO Zeba spin desalting column (Thermo Scientific), and incubated with azide-FIP-Cy5 at 4 °C overnight [59]. The dual-labelled antibody was purified using a 50K MWCO Amicon filter (Merck, Millipore), and the degree of labelling was measured by NanoDrop UV-vis spectrophotometer.

#### M.1.3 Internalisation assay

SHIP internalisation assays were performed by incubating the cells with dual-labelled anti-TFR antibody in DMEM containing 0.1% FBS at 37°C for different time points. After incubation, cells were washed thrice with cold PBS and resuspended in propidium iodide with or without quencher (1 μM), as described previously [59]. Cells were analysed using a Stratedigm S1000EON flow cytometer and FlowJo 10.8.0.

### M.2 Mathematical methods

#### M.2.1 Code availability

Codes are available on Github at https://github.com/ap-browning/internalisation.

#### M.2.2 Solution to dynamical model

The dynamical model is given by

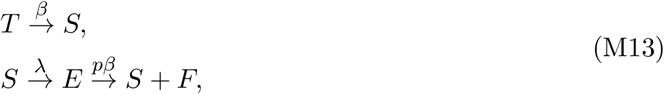

which we express as the system of ordinary differential equations

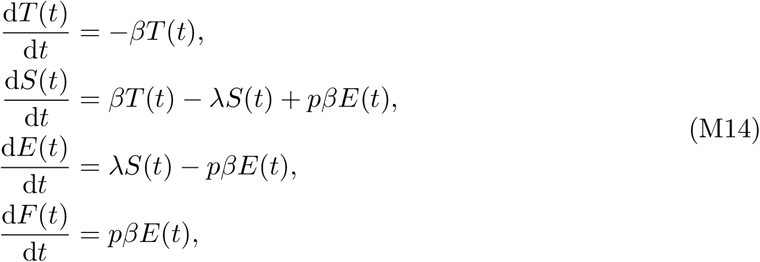

subject to the initial condition

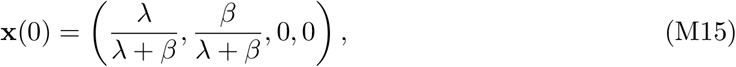

where **x**(*t*) = (*T*(*t*), *S*(*t*), *E*(*t*), *F*(*t*)). The solution to eq. (M14) is given by

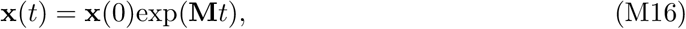

where

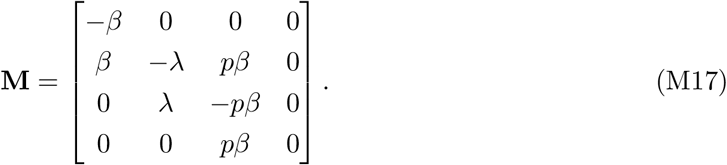

Equation (M16) is solved for *A*(*t*) = *S*(*t*) + *E*(*t*) + *F*(*t*) and *I*(*t*) = *E*(*t*) + *F*(*t*) exactly using Mathematica. Our implementation is given in `Module/Model/deterministic.jl`.

#### M.2.3 Inference using approximate Bayesian computation

We perform inference using approximate Bayesian computation (ABC) [38, 40]. Given a set of experimental observations 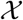, we encode knowledge about the model parameters ***θ*** in the *posterior distribution*, given by

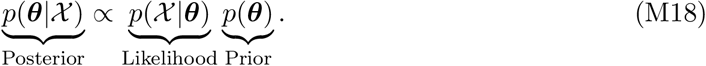

Here, *p*(***θ***) denotes the *prior distribution*, which encodes prior parameter knowledge. In our work, we take a standard approach and set the prior to be uniform with independent components [60]. We choose parameter bounds to reflect either physical constraints on parameters (i.e., all correlations are bounded and rates, standard deviations, proportionality constants are positive) or realistic bounds (for example, we expect the distributions of *λ* and *β* to be negatively skewed so that support is low, but non-zero, at zero if internalisation or recycling is inhibited in a small proportion of cells).

In ABC, we approximate the posterior distribution using the ABC posterior

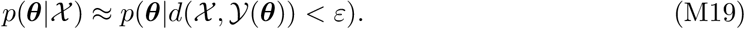

Here, 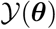 denotes synthetically generated observations of the model using parameters ***θ*** and *d*(·,·) is a *discrepancy measure* that measures how close synthetically generated observations lie to the experimental data. In ABC, *ε* is a parameter that describes the maximum discrepancy at which synthetic observations are judged to be close. In the following subsections, we outline our choice of discrepancy measure followed by our algorithm to sample from eq. (M19).

##### Discrepancy measure

We denote by 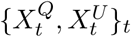 experimental observations from samples that are not quenched at time *t*, where *Q* and *U* denote measurements from the quenchable and unquenchable probes, 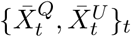 those of the quenched experiments, and 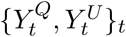 and 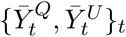 as *m* synthetically generate observations from the mathematical model. We denote the complete set of experimental observations 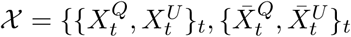 and similar for 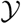.

Noting that the relationship between signals from the quenchable and unquenchable probes are generally linear, we develop a discrepancy metric, 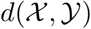 that matches univariate marginal distributions using a modified Anderson-Darling statistic *A*(*X, Y*), and matches the correlations between quenchable and unquenchable probes in the experimental and synthetic data. As we expect 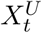 and 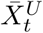 to be similar (since 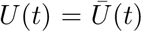), we apply a relative weighting of 1/2 to Anderson-Darling statistics involving the unquenchable probe.

Therefore,

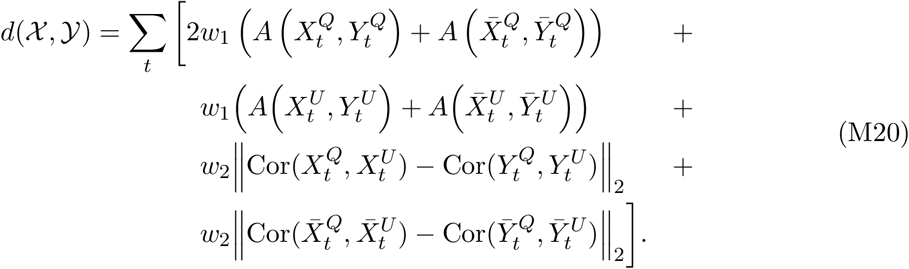

We set *m* = 1000, *w*_1_ = 1 and *w*_2_ = 80.

To compare experimental observations from a marginal distribution, 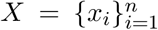, to a synthetically generated set of observations, 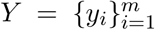, we employ a modified Anderson-Darling statistic. As the number of experimental observations, *n*, is large, we first compute an empirical distribution function *F*_obs_(*x*) using linear interpolation and *k* = 1000 equally spaced points from min_*i*_(*x_i_*) to max_*i*_(*x_i_*). The Anderson-Darling statistic is given by [44]

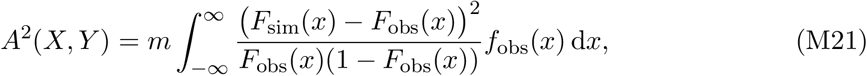

where *F*_sim_(*x*) is the empirical distribution function of the simulated data 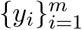.

Equation (M21) may be calculated by

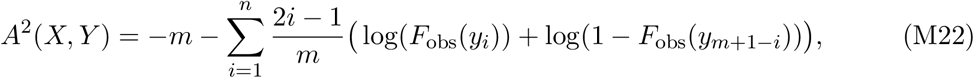

where 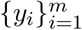 are sorted such that *y_i_* ≤ *y*_*i*+1_. To avoid blow up in cases where *y_i_* fall outside the range of the observed data 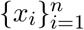 (where *F*_obs_(*x*) = 0 or 1) we set

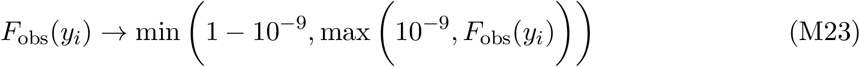

in eq. (M22). Our implementation is given in `Module/Inference/discrepancies.jl`.

##### Sampling algorithm

To identify regions of the parameter space with non-neglible posterior density, we employ the sequential Monte Carlo (SMC) algorithm described in [45]. In summary, we consider an initial sample of 1000 particles, which comprise parameter combinations sampled from the prior distribution with equal weight. Initially, a model realisation is produced for each particle and compared to the experimental data using eq. (M20). Particles are then sorted by the discrepancy metric and 750 particles with the lowest discrepancy are discarded. The ABC acceptance tolerance, *ε*, is set to be the largest discrepancy from the remaining particles. At each successive iteration, 750 replacement particles are resampled from the 250 weighted particles that remain and perturbed using a multivariate Gaussian perturbation kernel with covariance equal to twice the empirical covariance of the remaining 250 particles. This process is repeated until 750 replacement particles are found with a discrepancy lower than *ε*. Particles are reweighed according to the formula given in [45]. We iterate the algorithm until the acceptance rate at the resampling step is lower than the target 0.1%.

We treat the ABC SMC algorithm as a particle-based global optimisation algorithm, the result of which is the particle, 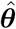, with the lowest discrepancy that is used to initiate and tune an ABC MCMC algorithm based on the Metropolis-Hastings algorithm [61] to explore the posterior. The ABC acceptance tolerance, *ε*, for the MCMC algorithm is chosen by taking the median discrepancy from 100 realisations of the model at 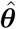. All ABC MCMC chains are then initiated at 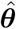.

We implement ABC MCMC using `MCMCChains.jl` [62]. First, we produce four pilot chains of length 10^6^ using a multivariate normal proposal with covariance equal to [63]

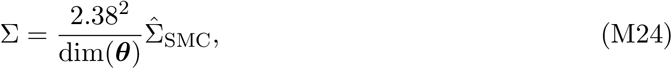

where 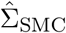 is the empirical covariance of the 1000 SMC particles. Every 100th sample is retained, and the covariance tuned using eq. (M24) where 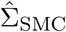 is replaced by the empirical covariance of the 40,000 samples identified using the pilot MCMC chains. Four tuned chains of length 10^7^ are produced and every 100th sample retained.

Our implementation of the SMC and MCMC algorithms are given in `Module/Inference/ abc.jl`. Our implementation of the ABC-SMC-MCMC sampling algorithm is given in `Results/ MainResult_Inference.jl`.

#### M.2.4 Dependence modelling with Gaussian copula

To produce realisations of the model we require independent samples of ***ξ***, a random variable described by marginal distributions where the dependence structure is described by a Gaussian copula. In this section, we outline a computationally efficiency algorithm to sample from the marginal distributions of ***ξ***. Additionally, we outline how we form an analytical expression for the bivariate marginal distributions used to produce visualisations in fig. 5.

##### Inverse transformation sampling

Sampling from the joint distribution *f*(*x, y, z*) with the dependence structure described by a Gaussian copula relies on the probability inverse transformation and, therefore, becomes prohibitively expensive if the quantile function of a marginal is expensive. Therefore, we employ an approximate sampling algorithm (algorithm 1) in cases where the quantile function of the marginal is expensive (i.e., when *x, y* or *z* are Gamma-distributed) and a large number of samples is required.

###### Algorithm 1 Probability inverse transformation sampling of *n* > 100 samples where quantile function *F*^−1^(*u*) is prohibitively expensive.

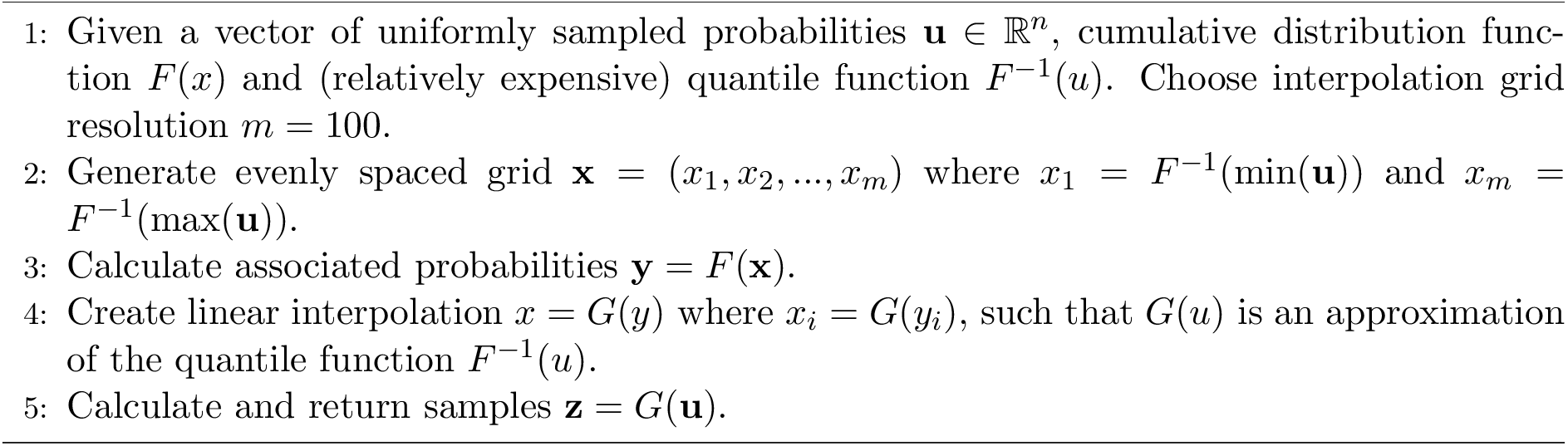

##### Bivariate marginal distributions

Consider the joint distribution of the random variables *X, Y*, and *Z*, with respective marginal density functions *f*(*x*), *f*(*y*) and *f*(*z*), distribution functions *u* = *F*(*x*), *v* = *F*(*y*) and *w* = *F*(*z*), connected through the Gaussian copula *C*(*u, v, w*). Then the marginal *f*(*x, y*) is given by

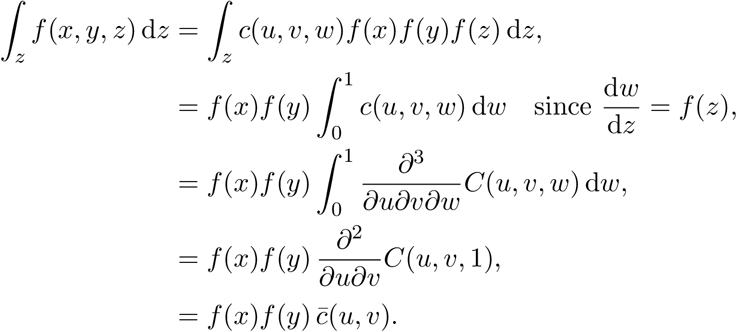

Here 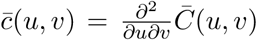, where 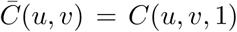 is the bivariate copula describing the marginalised dependence between *x* and *y*. Since *C*(*u, v, w*) is a Gaussian copula, we have that

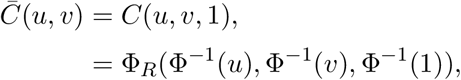

where Φ_Σ_ is the distribution function of the multivariate Gaussian distribution with covariance matrix Σ, and Φ(*x*) is the distribution function of the standard univariate Gaussian distribution. We also note that lim_*a*→∞_ Φ(*a*) = 1. Denoting *φ*_Σ_ the probability density function of the multivariate Gaussian distribution with covariance matrix Σ, we have that

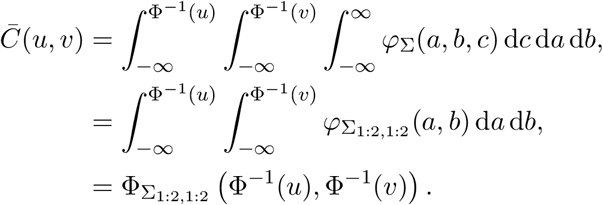

Therefore, 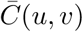 is a Gaussian copula with covariance matrix Σ_1:2,1:2_.

