## Supplementary text for "Identifying cell-to-cell variability in internalisation using flow cytometry"

### Supplementary material for "Identifying cell-to-cell variability in internalisation using flow cytometry"

#### Contents

**S1 Antibody-receptor binding** 2

**S2 Quenching efficiency** 3

**S3 Autofluorescence distribution** 4

**S4 Alternative parameterisation of Gamma distribution** 5

**S5 Parameters and MCMC diagnostics** 6

*These authors contributed equally.

### S1 Antibody-receptor binding

In the main text, we present a model where we assume that antibody binding occurs at a faster timescale to internalisation and recycling. Here, we include an additional compartment, $S_F$ that describes the number of free receptors on the cell surface. As the number of antibody molecules (approximately $2 \times 10^6$ per cell) is much larger than the overall number of receptors (approximately $4 \times 10^5$ per cell [1]) we assume that antibody binding occurs at a constant rate of $\gamma\,\text{min}^{-1}$. Therefore, the expanded model is given by

$$
\begin{aligned}
\frac{\mathrm{d}T}{\mathrm{d}t} &= -\beta T, \\
\frac{\mathrm{d}S_F}{\mathrm{d}t} &= \beta T + p\beta E - \gamma S_F, \\
\frac{\mathrm{d}S}{\mathrm{d}t} &= \gamma S_F - \lambda S, \\
\frac{\mathrm{d}E}{\mathrm{d}t} &= \lambda S - p\beta E, \\
\frac{\mathrm{d}F}{\mathrm{d}t} &= p\beta E,
\end{aligned}
\tag{S1}
$$

subject to the same equilibrium assumption as in the main document, so that

$$
\mathbf{x}(0) = \left( \frac{\lambda}{\beta + \lambda}, \frac{\beta}{\beta + \lambda}, 0, 0, 0 \right). \tag{S2}
$$

We perform profile likelihood analysis [2] to determine if $\gamma$ is identifiable from the data, results are shown in fig. S1. An approximate 95% confidence interval for $\gamma$ is given by $\gamma > 1.71$; i.e., $\gamma$ is one-sided identifiable, and is at least an order of magnitude larger than the parameters governing the internalisation and recycling rates.

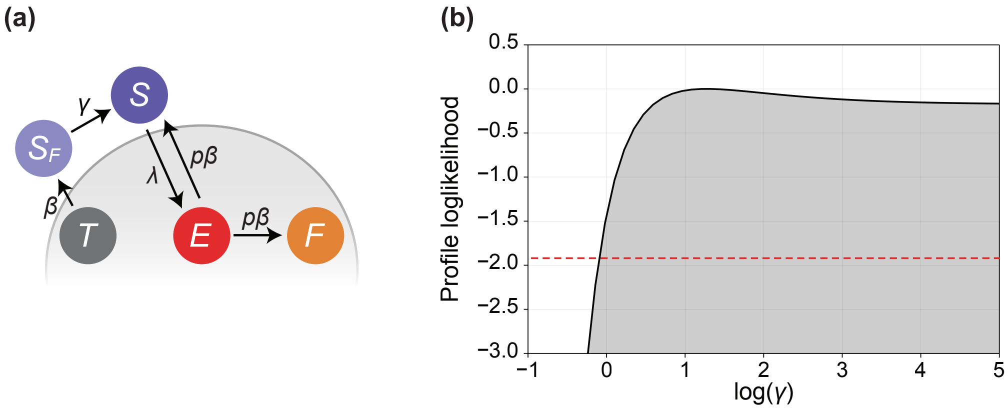


**Figure S1.** (a) The dynamical model describes the relative concentration of internal, transferrin bound receptors, $T$ (grey); surface bound receptors, $S_F$ (blue); surface antibody-bound receptors, $S$ (blue); internal antibody bound receptors, $E$ (red); and internal free antibody, $F$ (orange). (b) Profile log-likelihood for the antibody binding rate, $\gamma$. The red dashed line indicates the threshold for an approximate 95% confidence interval based on Wilks' theorem.

#### S2 Quenching efficiency

The *quenching efficiency*, denoted by $\eta$, describes the proportion of surface-bound fluorescent probes that are quenched through the introduction of the quencher dye. We pre-estimate the quenching efficiency using fluorescent measurements from samples kept at 4 °C, which inhibits internalisation.

*Standard approach*

Typically, the quenching efficiency is calculated based on the mean fluorescence intensity (MFI) of 4 °C samples, such that

$$\eta_{\mathrm{MFI}} = 1 - \frac{\overline{Q}^{4\,^{\circ}\mathrm{C}}_{\mathrm{MFI}}}{Q^{4\,^{\circ}\mathrm{C}}_{\mathrm{MFI}}}. \tag{S3}$$

Here, $Q^{4\,^{\circ}\mathrm{C}}_{\mathrm{MFI}}$ the MFI from the quenchable probe (i.e., Cy5 fluorophore-labelled probe) of 4 °C samples that have not been quenched. Similarly, $\overline{Q}^{4\,^{\circ}\mathrm{C}}_{\mathrm{MFI}}$ is that from samples that have been quenched. Clearly, for perfect quenching $\overline{Q}^{4\,^{\circ}\mathrm{C}}_{\mathrm{MFI}} = 0$ so that $\eta_{\mathrm{MFI}} = 1$.

We estimate

$$\eta_{\mathrm{MFI}} \approx 0.9398. \tag{S4}$$

We use the quenching efficiency calculated using MFI for analysis using the homogeneous model, which includes MFI measurements from data.

*Heterogeneous model*

For analysis using the heterogeneous model we calculate the quenching efficiency using the mean signal, such that

$$\eta = 1 - \frac{\langle \overline{Q}^{4\,^{\circ}\mathrm{C}} \rangle}{\langle Q^{4\,^{\circ}\mathrm{C}} \rangle}. \tag{S5}$$

Here, $\langle \cdot \rangle$ denotes the sample mean.

We estimate

$$\eta_{\mathrm{MFI}} \approx 0.9440, \tag{S6}$$

a slightly higher that the estimate based on MFI.

#### S3 Autofluorescence distribution

We build an empirical distribution for cellular autofluorescence, $(E_Q, E_U)$ using flow cytometry data from samples that are not introduced to labelled antibody (fig. S2).

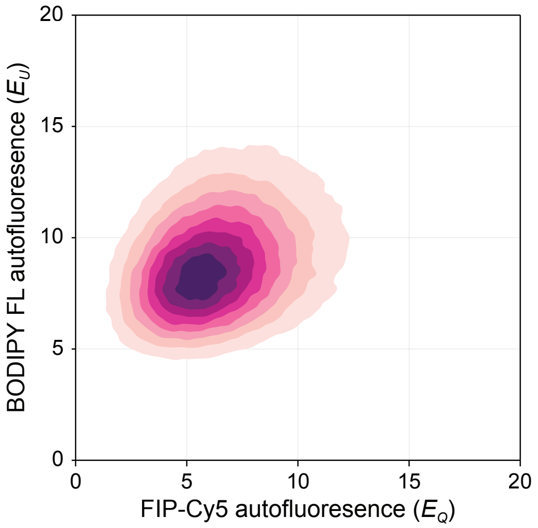


**Figure S2. Empirical autofluorescence distribution.** We obtain the autofluorescence distribution from samples that are incubated without antibody.

#### S4 Alternative parameterisation of Gamma distribution

To allow specification of the location, variance and skewness of the internalisation and recycling rates, we assume that $\lambda$ and $\beta$ are shifted Gamma variables. That is, we specify

$$a\tilde{\lambda} + b \sim \text{Gamma}(k, \theta), \tag{S7}$$

where $k$, $\theta$, $a$ and $b$ are chosen so that $\mathbb{E}(\tilde{\lambda}) = \mu_\lambda$, $\text{Std}(\tilde{\lambda}) = \sigma_\lambda$, and $\text{Skewness}(\tilde{\lambda}) = \omega_\lambda$. To ensure positivity, we assume

$$\lambda = \text{Truncated}(\tilde{\lambda}, 0, \infty), \tag{S8}$$

i.e., $\lambda$ is a truncated distribution that with only positive support. Similar assumptions are made for $\beta$.

In fig. S3 we fix $\mu_\lambda = 0.25$ and demonstrate various choices of $\omega$ and $\sigma$. As $\omega \to 0$, $\lambda$ approaches a normal distribution. For $\omega > 0$, $\lambda$ is positively-skewed, and for $\omega < 0$, $\lambda$ is negatively-skewed.

The alternatively parameterised Gamma distributions are implemented in `Module/Model/distributions.jl`.

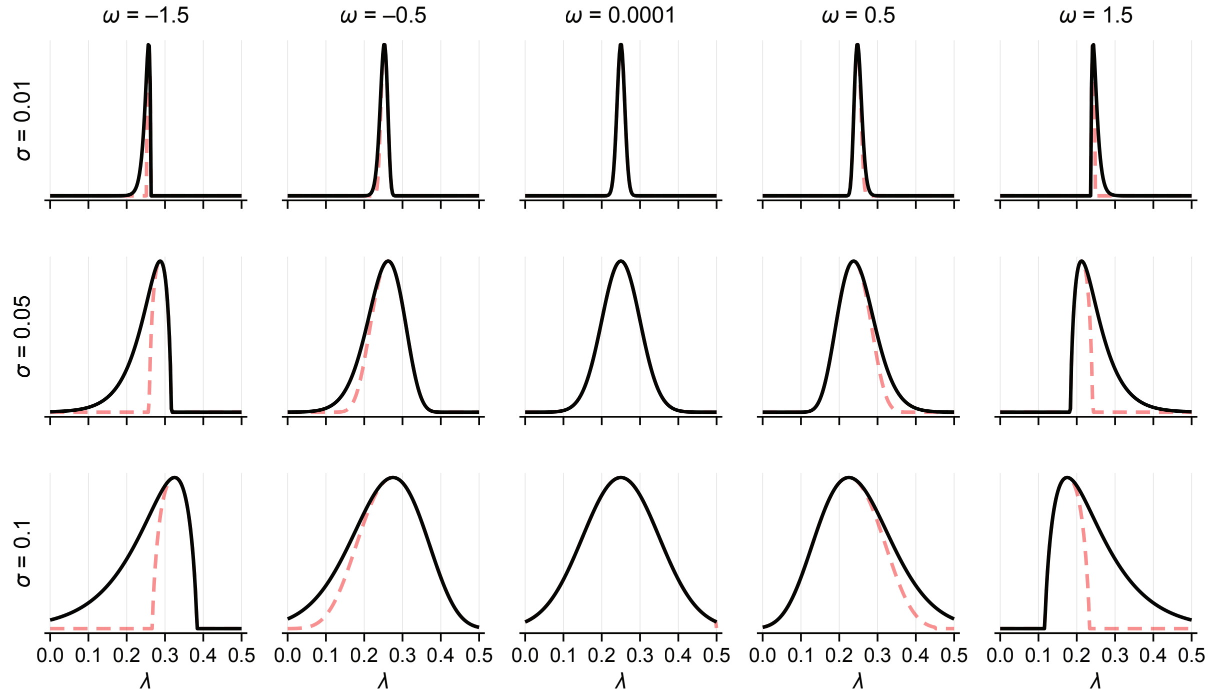


**Figure S3.** Probability density function for $\lambda_i \sim \text{ShiftedGamma}(\mu, \sigma, \omega)$ for various choices of spread, $\sigma$, and skewness, $\omega$. In all cases, we fix $\mu = 0.25$.

### S5 Parameters and MCMC diagnostics

| Parameter | | Best-fit | Mean | Std | ESS | $\hat{R}$ |
|---|---|---|---|---|---|---|
| $\alpha_1$ | Cy5-FIP intensity proportionality constant | 10130.0 | 9399.0 | 838.9 | 1629.0 | 1.021 |
| $\alpha_2$ | BDP-FL intensity proportionality constant | 47.39 | 43.71 | 3.964 | 1589.0 | 1.02 |
| $\sigma_1$ | Scaling factor for Cy5-FIP shot noise | 0.4338 | 3.618 | 2.112 | 2533.0 | 1.0 |
| $\sigma_2$ | Scaling factor for BDP-FL shot noise | 0.5638 | 0.2788 | 0.1549 | 2598.0 | 1.001 |
| $\mu_R$ | $R + c \sim \text{LogNormal}(\mu_R, \sigma_R), \quad \mathbb{E}(R) = 1$ | 0.127 | 0.2614 | 0.2085 | 1500.0 | 1.021 |
| $\sigma_R$ | | 0.386 | 0.3244 | 0.0789 | 1385.0 | 1.024 |
| $\mu_\lambda$ | Mean of $\lambda$ distribution | 0.1795 | 0.1163 | 0.04622 | 1966.0 | 1.006 |
| $\sigma_\lambda$ | Standard deviation of $\lambda$ distribution | 0.1108 | 0.1523 | 0.06189 | 1956.0 | 1.004 |
| $\omega_\lambda$ | Skewness of $\lambda$ distribution | 0.716 | −0.5245 | 0.8504 | 2284.0 | 1.004 |
| $\mu_\beta$ | Mean of $\beta$ distribution | 0.02944 | 0.04493 | 0.01701 | 1511.0 | 1.011 |
| $\sigma_\beta$ | Standard deviation of $\beta$ distribution | 0.0596 | 0.03938 | 0.02249 | 1210.0 | 1.028 |
| $\omega_\beta$ | Skewness of $\beta$ distribution | −0.1209 | −0.5606 | 0.8753 | 1938.0 | 1.006 |
| $\rho_{R\lambda}$ | Correlation between $R$ and $\lambda$ | −0.6416 | −0.1008 | 0.3686 | 1716.0 | 1.015 |
| $\rho_{R\beta}$ | Correlation between $R$ and $\beta$ | −0.6419 | −0.3468 | 0.4232 | 1024.0 | 1.064 |
| $\bar{\rho}_{\lambda\beta}$ | Scaled correlation between $\lambda$ and $\beta$ | 0.7101 | 0.2839 | 0.5157 | 1739.0 | 1.016 |
| $p$ | antiTDR/TFR disassociation probability | 0.04243 | 0.04796 | 0.008772 | 1780.0 | 1.013 |

**Table 1.** Model best-fit and MCMC statistics and diagnostics. Effective sample size (ESS) and $\hat{R}$ calculated using `MCMCChains.jl` [3] based on [4]
.
